# High fat diet induces differential age- and gender-dependent changes in neuronal function linked to redox stress

**DOI:** 10.1101/2024.07.12.603123

**Authors:** Megan de Lange, Vladyslava Yarosh, Kevin Farell, Caitlin Oates, Renee Patil, Isabel Hawthorn, Mok-Min Jung, Sophie Wenje, Joern R Steinert

## Abstract

The prevalence of neurodegenerative diseases, such as Alzheimer’s disease and Parkinson’s disease, is steadily increasing, posing significant challenges to global healthcare systems. Emerging evidence suggests that dietary habits, particularly consumption of high-fat diets specify which fats, may play a pivotal role in the development and progression of neurodegenerative disorders. Moreover, several studies have shed light on the intricate communication between the gut and the brain, known as the gut-brain axis and its involvement in neurodegenerative processes.

This study aims to assess the effects of a high-fat dietary intake on various aspects of neuronal function during aging and following gender separation to help understand the potential contributions of diet to neuronal function.

To investigate the effects of a high-fat diet, *Drosophila melanogaster* was used and exposed to standard normal food diet (NF) and high-fat diet (HF). Adults were grouped at 10 and 45 days of age in male and female flies reared under the same conditions. Multiple assays were conducted, showing differential gender- and HF diet-induced oxidative stress levels as determined by malondialdehyde (MDA) measurements, enhanced caspase-3 expression and reduced climbing activity. Adult lifespan under both dietary conditions was unchanged but odour-associated learning ability was reduced in larvae reared in a HF diet.

This is the first study to characterise effects of diet on neuronal phenotypes in an age- and gender-specific manner in a *Drosophila* model. Our findings suggest a HF diet induces differential forms of neuronal dysfunction with age and sex-specific outcomes, characterised by enhanced oxidative stress and cell death which impacts on neuronal and behavioural functions.

## Introduction

Various neurodegenerative diseases are characterized by progressive loss of neuronal function and structure, leading to cognitive decline or motor dysfunction and ultimately, disability and death. While aging is a major risk factor for neurodegeneration (Azam, Haque et al. 2021), growing evidence implicates environmental factors, including diet, in various pathologies (Popa-Wagner, Dumitrascu et al. 2020). In particular, high-fat diets, typical of Western dietary patterns, have been linked to increased risk of neurodegenerative disorders (Graham, Harder et al. 2016, Wieckowska-Gacek, Mietelska-Porowska et al. 2021, Liang, Gong et al. 2023, Valentin-Escalera, Leclerc et al. 2024). Understanding the molecular mechanisms underlying the effects of high-fat diets in neurodegeneration is crucial for developing preventive and therapeutic strategies.

High-fat diets (HFD), rich in saturated fats and cholesterol, have been shown to induce chronic low-grade inflammation in various tissues, including the brain (Graham, Harder et al. 2016, Spencer, D’Angelo et al. 2017, Gonzalez Olmo, Bettes et al. 2023). Resulting neuroinflammation and redox stress play central roles in the pathogenesis of various neurodegenerative diseases by promoting neuronal damage and impairing synaptic function as shown in different model systems (Bourgognon, Spiers et al. 2021, Spiers, Cortina Chen et al. 2022, Steinert and Amal 2023, Zhang, Xiao et al. 2023, Adamu, Li et al. 2024). Thus, a consumption of high-fat diets leads to activation of microglia, the resident immune cells of the brain, and release of pro-inflammatory cytokines and reactive oxygen species, contributing to neuroinflammatory responses observed in neurodegenerative conditions.

In addition, the gut microbiota, composed of trillions of microorganisms inhabiting the gastrointestinal tract, plays a critical role in maintaining gut homeostasis and modulating host physiology (Sommer and Backhed 2013, Hou, Wu et al. 2022). Disruption of the gut microbiota composition, known as dysbiosis, has been associated with various diseases, including neurodegeneration (Hou, Wu et al. 2022, Intili, Paladino et al. 2023, Solanki, Karande et al. 2023). High-fat diets can alter the gut microbiota composition, favoring the growth of pro-inflammatory bacteria and reducing microbial diversity (Deshpande, Saxena et al. 2019, Hamamah, Amin et al. 2023). The resulting dysbiosis-induced alterations in gut permeability and integrity facilitate the translocation of microbial products, metabolites and inflammatory mediators into the systemic circulation, ultimately impacting brain function and facilitating neurodegeneration (Di Vincenzo, Del Gaudio et al. 2024).

Previous reports suggest that the intake of HFD increases the risk of obesity and type 2 diabetes in approximately 9% of the human population (Teodoro, Varela et al. 2014) causing oxidative stress, cellular inflammatory response and impacts on cognitive function as assessed in animal studies (Tan and Norhaizan 2019). Extensive research has shown that diet-induced obesity potentially leads to memory impairment in rodents (Alzoubi, Mayyas et al. 2018). Mice fed a long-term HFD show reduced learning and memory performance (Cordner and Tamashiro 2015, Hsu, Sheen et al. 2020), as well as depressive- (Kurhe, Mahesh et al. 2014) and anxiety-like behaviors (Sivanathan, Thavartnam et al. 2015, Zemdegs, Quesseveur et al. 2016). Although most studies have been performed in mixed or single gender test groups, some data point towards gender-specific effects of diet (Murtaj, Penati et al. 2022, Stapleton, Welch et al. 2024), whereas other studies could not confirm sex-specific dietary effects on neuronal function (Underwood and Thompson 2016).

In addition to rodent models, *Drosophila* has emerged as an important model system to assess dietary impacts on neuronal function (Huang, Song et al. 2020, Alassaf and Rajan 2023) and fundamental metabolic regulatory processes in many disorders including obesity and diabetes (Alfa and Kim 2016). Previous studies have used approaches to study the effects of a HFD on various aspects of *Drosophila* physiology. As such, a 2% HFD (virgin coconut oil) induced developmental delay, growth defects and sleep fragmentation (Nayak and Mishra 2021). There is convincing evidence that a HFD reduces life span in *Drosophila* when exposed to between 20% to 30% coconut oil or 10% lard (Trindade de Paula, Poetini Silva et al. 2016, Wang, Sun et al. 2017, Wen, Zheng et al. 2018, Rivera, McHan et al. 2019, Chattopadhyay and Thirumurugan 2020, Liao, Amcoff et al. 2021, Nayak and Mishra 2021), however, due to the variable dietary and rearing conditions an interpretation of data can be challenging.

To investigate potential effects of a high fat diet on neuronal function and a gender specificity of effects, we exposed male and female *Drosophila* to a diet containing 10% virgin coconut oil for up to 40 days and assessed in 3^rd^ instar larvae and adult flies, at both sexes, pathology markers like oxidative stress and apoptosis as well as behavior like learning and locomotion.

## Methods

### Drosophila husbandry and maintenance

*Drosophila melanogaster* (w^1118^) in the NF experimental group were reared on a standard cornmeal-agar media, without active yeast, made up of water, cornmeal, agar, soya flour, yeast, sugar syrup, propionic acid and nipagin. *Drosophila melanogaster* in the high fat diet (HF) experimental group were reared on the same media to which organic virgin coconut oil (10% w/w) was added at day 5. All adult flies (males and unmated females) were tipped into new food vials every other day. Larvae were reared at the HF from day 0. All flies were kept at 18°C and a 12 h light-dark cycle.

For each experimental condition, adult heads were collected at 10 and 45 days of age, placed into a 1.5 ml microcentrifuge tube and stored at −80°C. Larvae were used at 3^rd^ instar developmental stage.

### MDA Assay

For each condition, 16 adult heads were placed into lysis buffer, homogenised (Kontes, Pellet Pestle Motor) for five minutes and centrifuged (MiniSpin Plus, Eppendorf) for 10 min at 12000 rpm. A standard curve for the MDA Assay was generated and MDA was measured using a kit in accordance with the manufacturer’s instructions (SIGMA-ALDRICH Lipid Peroxidation malondialdehyde (MDA) Assay, Catalog no. MAK085). In detail, a thiobarbituric acid (TBA) solution was prepared and stored at 4°C. To create the samples, 200 µl of the lysate was pipetted into a separate 1.5 ml microcentrifuge tube. An MDA-TBA adduct was then created by adding 600 µl of the TBA solution into each tube containing the standards and the samples. The tubes were incubated at 95°C for 60 minutes. After incubation, the tubes were cooled to room temperature for 10 minutes. 200 µl was then pipetted into a clear-bottomed 96-well microplate for analysis, with each sample and standard being repeated in triplicate. The absorbance was measured at 532 nm using the Clariostar and normalized data are expressed following the Bradford assay protein calculations.

### Immunohistochemistry

Following paraformaldehyde (PFA 4%) fixation, brains were processed in blocking and antibody dilution buffers (1x PBS/ 5% normal serum [NGS]/ 0.3% Triton X-100) for 60 minutes. Primary antibody (Cell Signalling Technology, rabbit anti-cleaved Caspase-3 antibody (Asp175)) was added (1:400) and incubated overnight at 4°C. The tubes were then placed on a rocker for 10 minutes to wash the brain in each tube and washed 3 times in PBS. Following the last wash, the secondary antibody (Invitrogen, Alexa Fluor 647 goat anti-rabbit antibody IgG) was added and samples incubated for 1-2 hours at room temperature in the dark.

The secondary antibody was removed, and samples incubated with Prolong Gold Antifade Reagent with DAPI (Sigma, Fluoroshield with DAPI).

### Confocal imaging

Brains were imaged using a confocal microscope (Zeiss LSM 880). The quantification of neurons undergoing apoptosis was done by assessing the extent of immunofluorescence intensity (cleaved caspase-3 antibody). Z-stacked images of the full brain at 20x magnification were initially captured. Additionally, the mid-brain region was focused on and Z-stacked images at 40x magnification were captured at 365nm (DAPI) and 633nm (Alexa Fluor 647) excitation. All images were quantified following constructions of maximal intensity projections in ImageJ v 1.53 (DAPI and cleaved caspase-3).

### Longevity assay

A minimum of 30 flies was separated at 5 days into their respective experimental groups to give different experimental conditions according to age, sex and diet. This set up was repeated to give a minimum of 4 vials per experimental condition. Flies were tipped into new vials every other day and kept at 18°C. The number of flies in each vial was counted every day or every second day and was recorded until all flies were dead to generate life span curves.

### Climbing assay

A climbing assay protocol was created to observe mobility and motor function. The testing chamber for the assay was composed of a vial, with markings at five centimeters and eight centimeters, and a sponge stopper. The vials were labelled according to the experimental group of flies in the vial. Polystyrene shields were used to block off natural light from the sides of the tubes. Ten flies were transferred to the testing chamber vials, using carbon dioxide anaesthesia, and were left to rest for 30 minutes to minimise the effect of anaesthesia on climbing ability. Following this resting period, the flies were tapped to the bottom of the vial and the number of flies above the eight-centimeter mark after eight seconds was recorded. The flies were allowed to recover for an additional minute and then the assay was repeated. This was done three times to generate three different recordings.

### Odour-associative learning

The odour-associated learning paradigm was used as reported previously (Saumweber, Husse et al. 2011, Stone, Cujic et al. 2023). Innate preferences were tested first by placing age-matched 3^rd^ instar larvae in a petri dish with the experimental odour on one side, leaving larvae to crawl for 3 minutes and recording larval position on the dish. For paired training, cohorts of 10-15 larvae were placed at the centre of a Petri dish (9 cm inner diameter) filled with 1% agarose, supplemented with fructose (2 M) as a taste reward (+) which included an odour-containing filter paper (n-amylacetate (AM), diluted 1:20 in paraffin oil). Paraffin has no behavioural significance as an odour (Saumweber, Husse et al. 2011). For training, larvae were placed at the midline and free to move for 3 min at a dish containing AM filter paper at each of the opposing edges of the Petri dish and fructose-containing agar (AM+). Subsequently, they were transferred to a Petri dish without fructose which had two non-AM filter papers (EM), and they were left there for the same amount of time. After such AM+/EM training sessions, they were transferred to the centre of a test Petri dish, where an AM odour filter paper was presented on one side and were thus tested for their preference for AM. After 3 min, the numbers of larvae (#) on the AM side, on the EM side, and in a 10-mm wide middle zone were counted. Larvae crawling up the sidewalls of the Petri dish were counted for the respective side, whereas larvae on the lid were excluded from the analysis. A preference (Pref) was calculated:

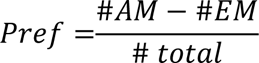

Preference indices range from +1 to −1, with positive values indicating preference and negative values indicating avoidance of AM. Across repetitions of the experiments, in half of the cases the sequence was as indicated (AM+/EM), whereas in the other cases it was reversed (EM/AM+). The procedure for unpaired training was the same, except that the Petri dishes featured either only AM or only the reward. After such AM/EM+ training (again in half of the cases the sequence was reversed: EM+/AM), the preference test was carried out as above. From the Pref scores after paired and unpaired training, a Preference Index (PI) was calculated:

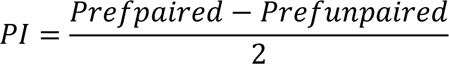

Performance indices range from +1 to −1. Positive PIs indicate appetitive associative memory, whereas negative values indicate aversive associative memory.

### Locomotor Activity

Age-matched third instar larvae (∼120 h of age) were selected, washed and placed onto a moist, food-free surface at a constant temperature of ∼20°C. Crawling activities were imaged over 10 min using AnyMaze software v7.16 (Stoelting Co., Wood Dale, IL, USA) and data was analysed off-line as reported previously (Robinson, Nugent et al. 2014).

### Statistical Analysis

Statistical analyses of the data were conducted using GraphPad Prism v10 (GraphPad Software Inc., San Diego, CA, USA). A two-way ANOVA or one-way ANOVA test was used when applicable with a posterior test (with Tukey’s multiple comparison). After confirming normal data distributions (Shapiro-Wilk test) and homogeneity of variance (Bartlett’s test), parametric or non-parametric statistics were applied. Unpaired Student’s t-test was used to compare locomotor and immunofluorescence data. Cumulative survival curves are presented and compared using the log-rank (Mantel-Cox) test. To compare PI and Pref AM data against a chance level (0), a one-sample t-test or Wilcoxon signed-rank test (odour learning) was used. Data are expressed as mean ± SEM or box and whisker blots where n is the number of flies and brains as indicated. Significance is shown as *p/# < 0.05, **p < 0.01, ***p < 0.001 and ****p < 0.0001.

## Results

### Larval locomotor activity and learning and memory

We wanted to assess how a HF diet impacts on neuronal function of the central nervous system (CNS) including the ventral nerve cord (VNC) of *Drosophila* larvae by characterizing odour-associated learning and locomotor activities. Initial experiments assessed crawling activity as a measure of locomotion that occurs as a result of the peristaltic contractions of ∼30 muscles (Landgraf, Bossing et al. 1997) associated with body segments (Heckscher, Lockery et al. 2012). Following tracking of a 10 min crawling period on a moist food-free surface, we quantified travel distance for control NF and HF diet fed larvae. Both groups travelled similar distances (p>0.05, Student’s t-test) indicating that the neuronal circuitries responsible for mediating locomotion are not affected by a HF diet (**Figure 1**).

**Figure 1:**
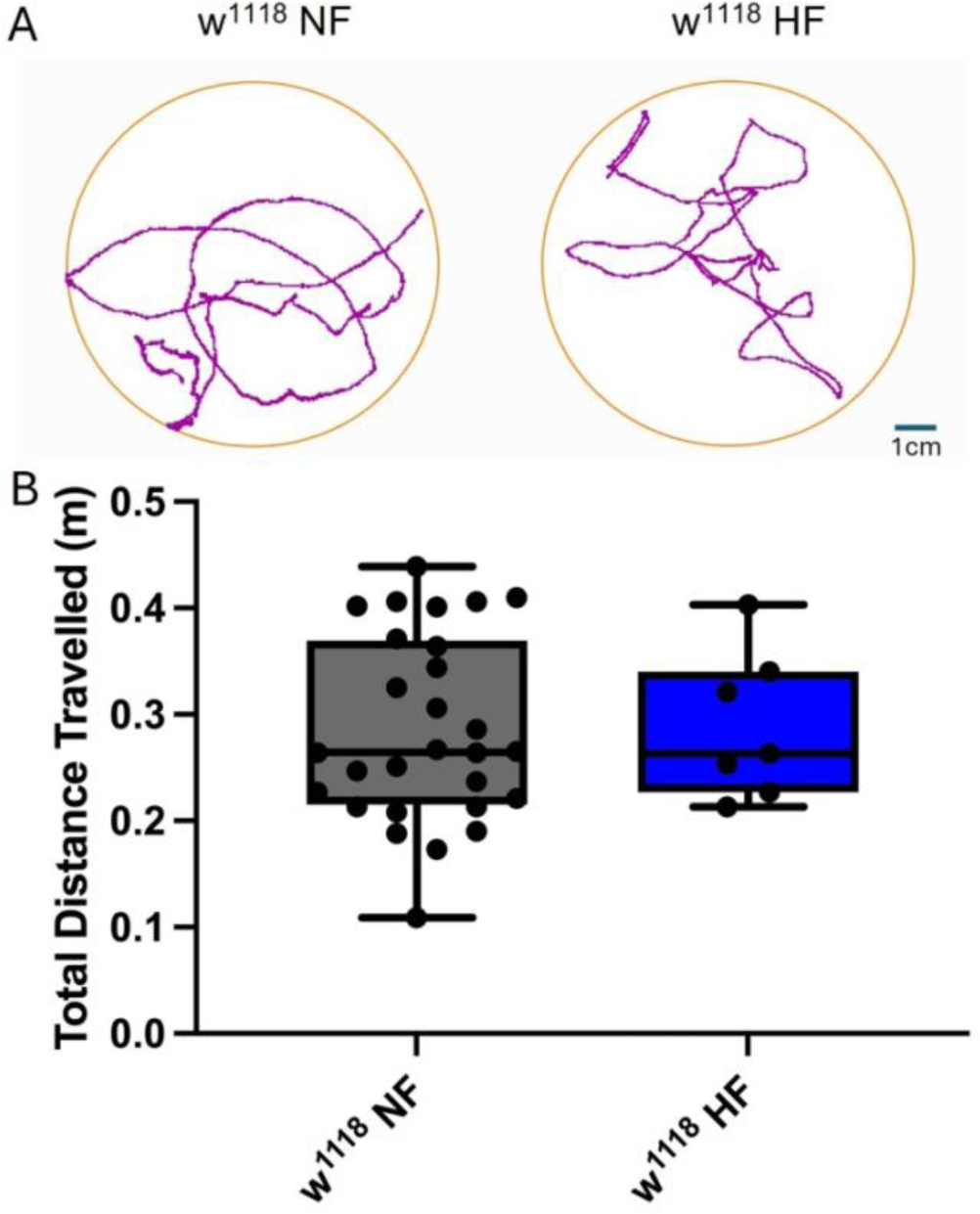
Locomotor activity is not affected by a HF diet. A, Larval crawling activities were recorded over 10 min within a crawling arena and tracks were recorded and plotted using AnyMaze software. B, Average crawling activities show no differences between both groups (n=7-29 larvae per condition, p>0.05, unpaired Student’s t-test).

A further CNS-driven and more complex behavior underlies learning and memory function in larvae which is highly plastic and adapts to experience (review in (Thum and Gerber 2019)). Mushroom Bodies (MB) in the larval brain are necessary for the formation of associative memory (Pauls, Selcho et al. 2010). This memory is expressed by changing navigation towards a cue (e.g. an odour) that has been associated with a reward (e.g. sucrose) or a punishment (e.g. quinine, electric shock) and can be assessed in odour-associated learning and memory experiments. In contrast to the lack of diet effects on locomotor activity, our data show that learning and memory function was affected by a HF diet as shown by reduced PI values of the HF diet test group (**Figure 2A**). As larvae also exhibit innate preference to various odours (Saumweber, Husse et al. 2011), we confirmed that the innate preference to AM was not affected by diet (**Figure 2B**).

**Figure 2:**
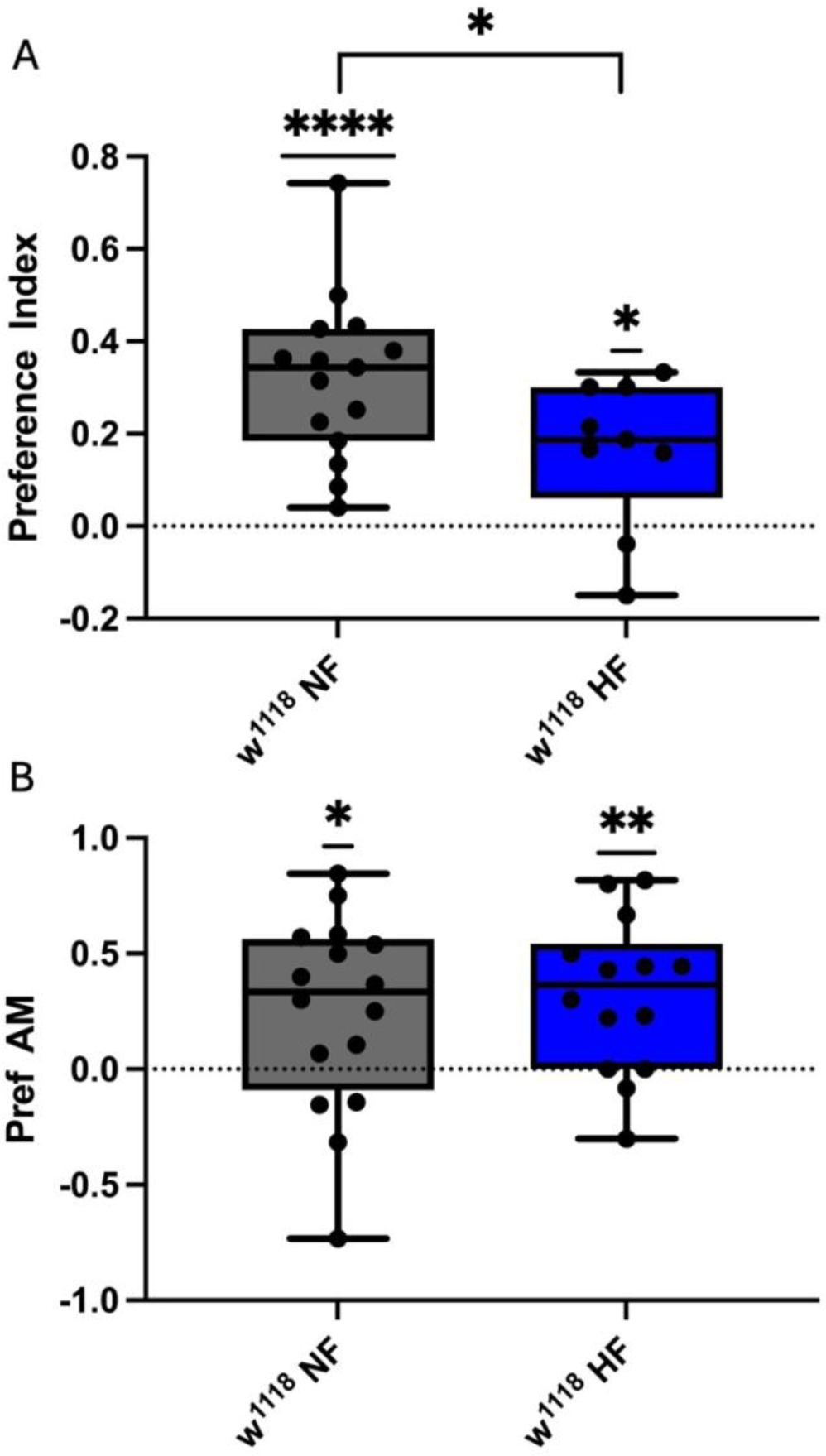
A HF diet reduces larval learning and memory function. A, Preference index (PI) was calculated for 3rd instar Drosophila larvae reared at both food conditions. PI values were significantly reduced in the HF diet group compared to the control group (Mann-Whitney test: NF vs HF: p=0.0393). PI vales for both groups were significantly different from zero (one-sample t-test, NF: ****p<0.0001, HF: *p=0.0161). B, Innate preferences to AM did not differ between control and diet groups, however, both groups exhibited a similar innate attraction to AM (one-sample t-test: NF: *p=0.034, HF: **p=0.0033, Student’s t-test: NF vs HF: p=0.6033, n=28-40).

### Negative geotaxis is affected by a HF diet

As the above reported data in larvae confirm neuronal changes after relatively short-term exposures to a HF diet, a developmental impact cannot be excluded, therefore we next assessed adult flies exposed to different diets from the age of 5 days. This excludes major developmental effects and allows testing for impacts of diets on the aging process (Li and Hidalgo 2020). A well-established readout of aging processes is performance of negative geotaxis-driven climbing activity (Rhodenizer, Martin et al. 2008, Piper and Partridge 2018). We tested male and female flies at the age of 10 days and 45 days (exposed to the HF diet for 5 and 40 days, respectively) and determined the climbing index. Aged flies showed a reduction in climbing abilities irrespective of diet and gender (**Figure 3**) as previously observed in mixed gender populations under standard food conditions (Rhodenizer, Martin et al. 2008). Importantly, we observed a negative effect of a HF diet at 10 days only in female flies. Control female flies had higher climbing indices than control male flies at 10 days (diet: F (3, 36) = 7.304, p=0.0006, age: F (1, 36) = 39.82, p<0.0001, two-way ANOVA, **Figure 3**), illustrating gender differences in climbing under control diets and gender-specific sensitivities to a HF diet.

**Figure 3:**
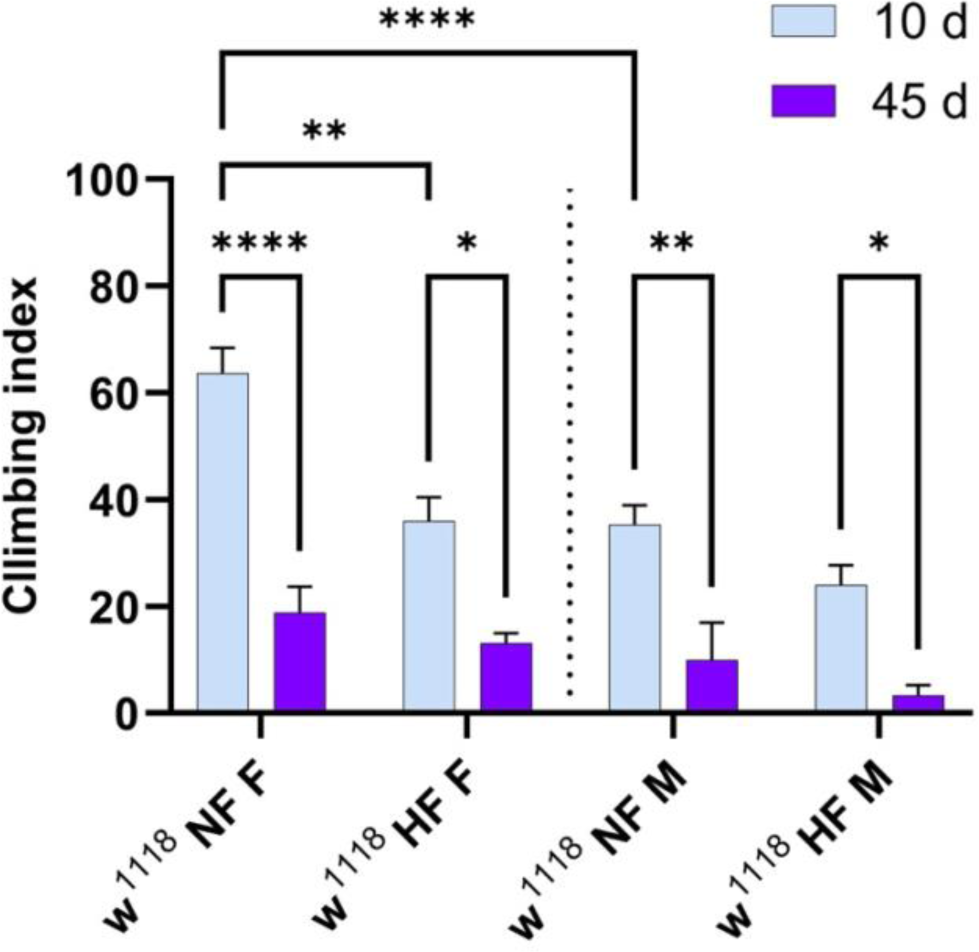
Diet and age impact on climbing abilities. A climbing assay was performed to assess motor function in both groups which had been fed either a NF or HF diet. The assay was performed at 10 or 45 days (d) of age. The data showed significant age- and gender-dependent changes (diet: F (3, 36) = 7.304, ***p=0.0006, age: F (1, 36) = 39.82, ****p<0.0001 (two-way ANOVA, n=10 flies per vial).

### HF diet does not alter longevity at lower temperature

Having observed different effects on neuronal function following a HF diet exposure, we next assessed adult survival under the same food conditions. A HF diet (2%, 10% and 20% coconut oil) has been shown previously to reduce life span when flies were reared at ∼25°C (Trindade de Paula, Poetini Silva et al. 2016, Nayak and Mishra 2021). Since flies can get trapped in fluid coconut oil residues at ∼25°C, we performed a longevity assay to observe diet-dependent effects on lifespan at 18°C rearing temperature which provided a solidified coconut oil-food mixture. Survival and neurodegeneration show inverse relationships with rearing temperature with higher temperatures accelerating cell death rates and reducing life span (Miquel, Lundgren et al. 1976, Molon, Dampc et al. 2020). We determined survival in male and female flies under both diets and analyzed data using a log-rank test. As expected, all life spans were prolonged at 18°C (Molon, Dampc et al. 2020) compared to studies performed at ∼25°C, however, neither diets nor gender had any effects on survival (**Figure 4**, log-rank (Mantel-Cox) test, p>0.05) with median survival rates for each condition showing no differences.

**Figure 4:**
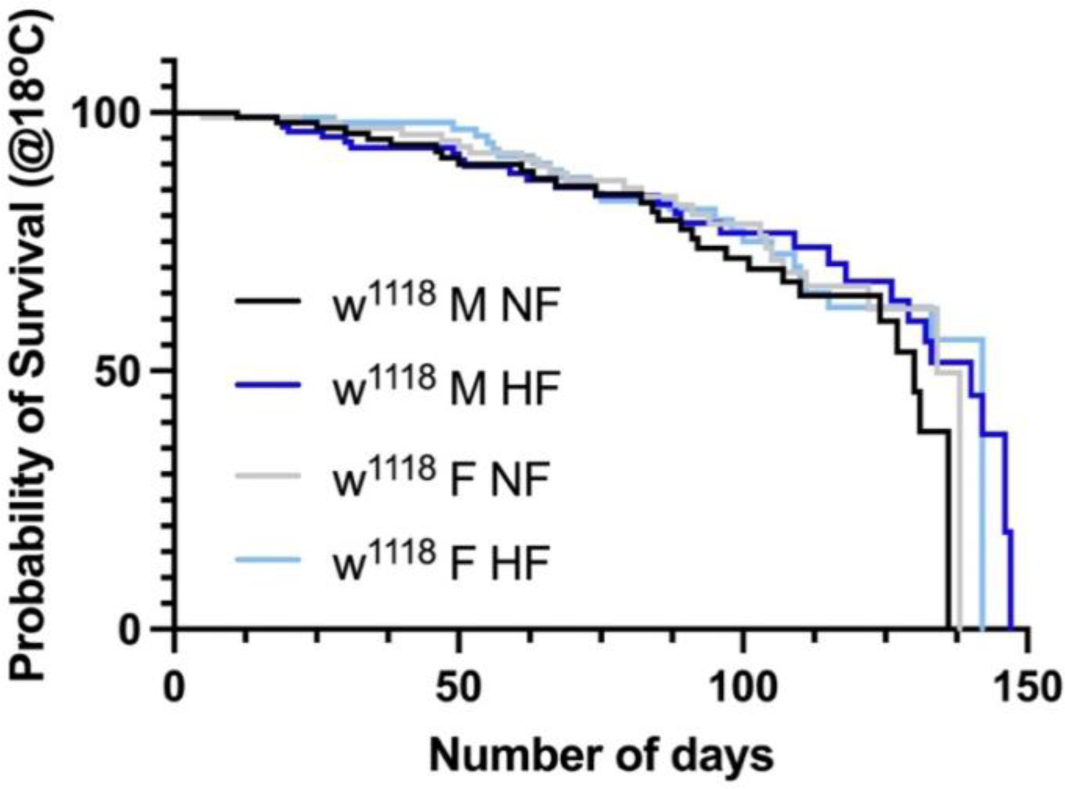
Longevity is not affected by a HF diet at 18°C rearing temperature. Survival curves for 4 conditions, log-rank (Mantel-Cox) test, p>0.05. Male NF vs Male HF: p=0.303 (median survival 130 vs 140 days), Female NF vs Female HF: p=0.764 (median survival 134 vs 142 days), Male NF vs Female NF: p=0.323 (median survival 130 vs 134 days), Male HF vs Female HF: p=0.9154 (median survival 140 vs 142 days), n=30-40 flies, minimum of 4 vials per experiment.

### Oxidative stress is increased following a HF diet

As aging and neurodegeneration and a high fat dietary intake have been associated with higher levels of cellular oxidative stress (Pratico 2002, Ayala, Munoz et al. 2014, Trindade de Paula, Poetini Silva et al. 2016, Nayak and Mishra 2021, Gomes, Dos Santos et al. 2023, Steinert and Amal 2023) which could be responsible for the observed behavioural changes and neuronal damage, we next assessed lipid peroxidation in fly brains by measuring levels of the secondary product during lipid peroxidation, malondialdehyde (MDA) (Ayala, Munoz et al. 2014) at both sexes and time points. Figure 5 shows quantifications of MDA levels for each experimental group. A two-way ANOVA comparison revealed that the levels of MDA were altered in a diet-dependant manner (diet: F (3, 16) = 33.41, p<0.0001 and age: F (1, 16) = 0.02156, p=0.8851, two-way ANOVA, **Figure 5**). MDA levels increased with age at a normal diet in both sexes (male: p=0.00182, female: p=0.0240), however, a HF diet is paralleled by a reduction in MDA levels at 45 days in both sexes compared to 10 days (male: p<0.0001, female: p<0.0001). A HF diet alone caused higher MDA levels measured at 10 days in both sexes (male: p<0.0001, female: p=0.0293). Interestingly, at a control NF diet, male flies show lower

**Figure 5:**
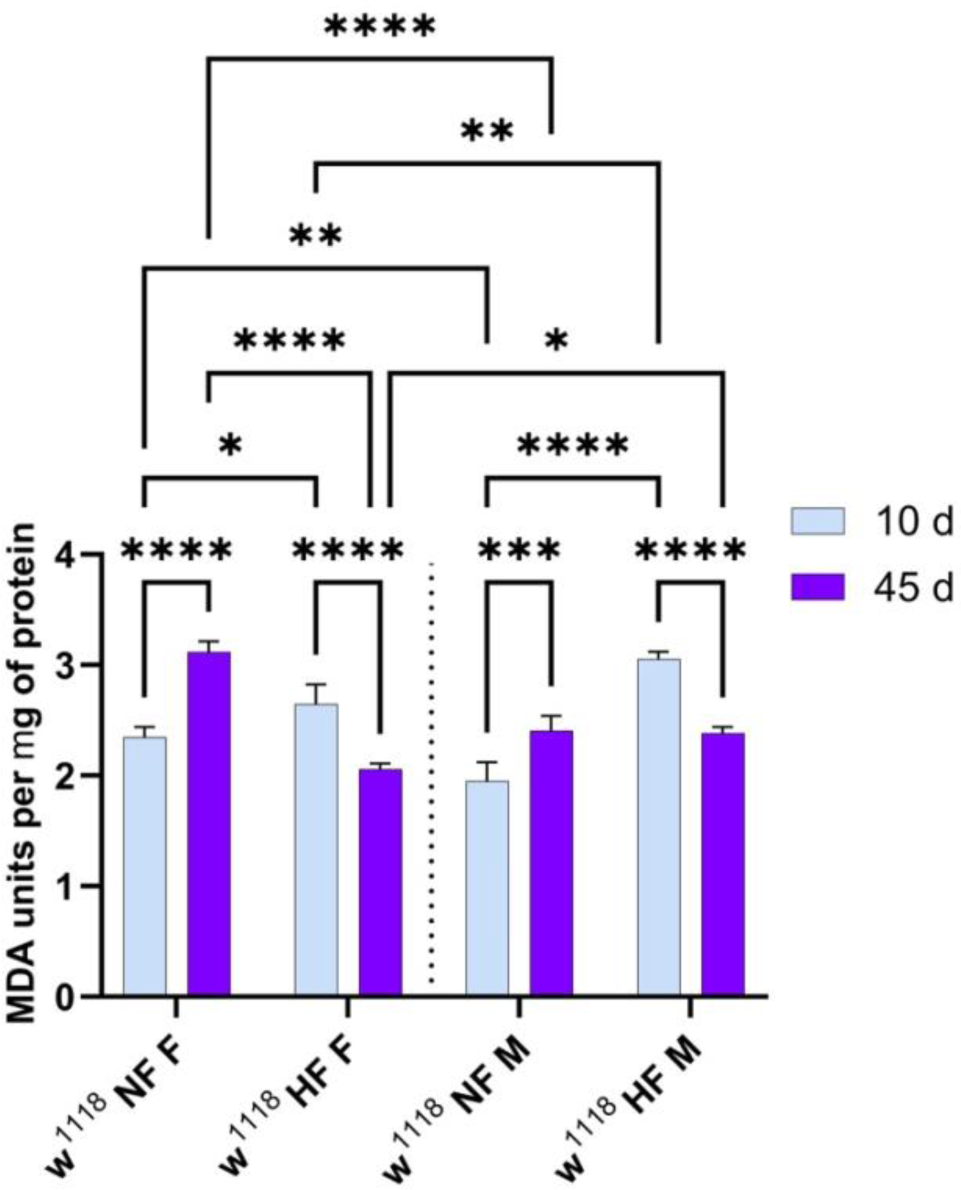
Age- and gender-specific changes in MDA levels following a HF diet. Levels of MDA were assessed expressed as arbitrary units per mg of protein. Age- and diet-dependent changes are shown for 10 and 45 days (d) (diet: F (3, 16) = 33.41, ****p<0.0001, age: F (1, 16) = 0.02156, p=0.8851, two-way ANOVA, n=16 heads per condition).

MDA levels at 10 days (p=0.0036). This data suggests an overall increase in MDA levels at 10 days following a HF diet but a reduction in MDA levels after 45 days of a HF diet in both sexes and highlights further gender-specific differences under both diets.

### Caspase-3 expression is elevated following HF diet exposure

As enhanced lipid peroxidation is causally involved in cell death and apoptotic pathway regulation (Ayala, Munoz et al. 2014), we next tested whether apoptosis was enhanced under the conditions of a HF diet. Immunostaining of *Drosophila* brains was performed to detect levels of apoptotic cell death in the CNS by analysing z-projections of maximal intensities of the caspase-3 labelling. This staining detects *Drosophila* DRONC (*Drosophila* Nedd2-like caspase) activity indicative of cleaved caspase-3 signalling (Fan and Bergmann 2010, Fan, Lee et al. 2010, Klepsatel, Galikova et al. 2016). We could not confirm gender differences at any tested condition and decided to merge sexes for the following analyses. Using a two-way ANOVA we only detected an increase in caspase signal at 45 days of age following a HF diet exposure (diet: F (1, 14) = 12.88, p=0.0030, age: F (1, 14) = 3.688, p=0.0754; NF: 45 days vs HF: 45 days, p=0.0093, **Figure 6**).

**Figure 6:**
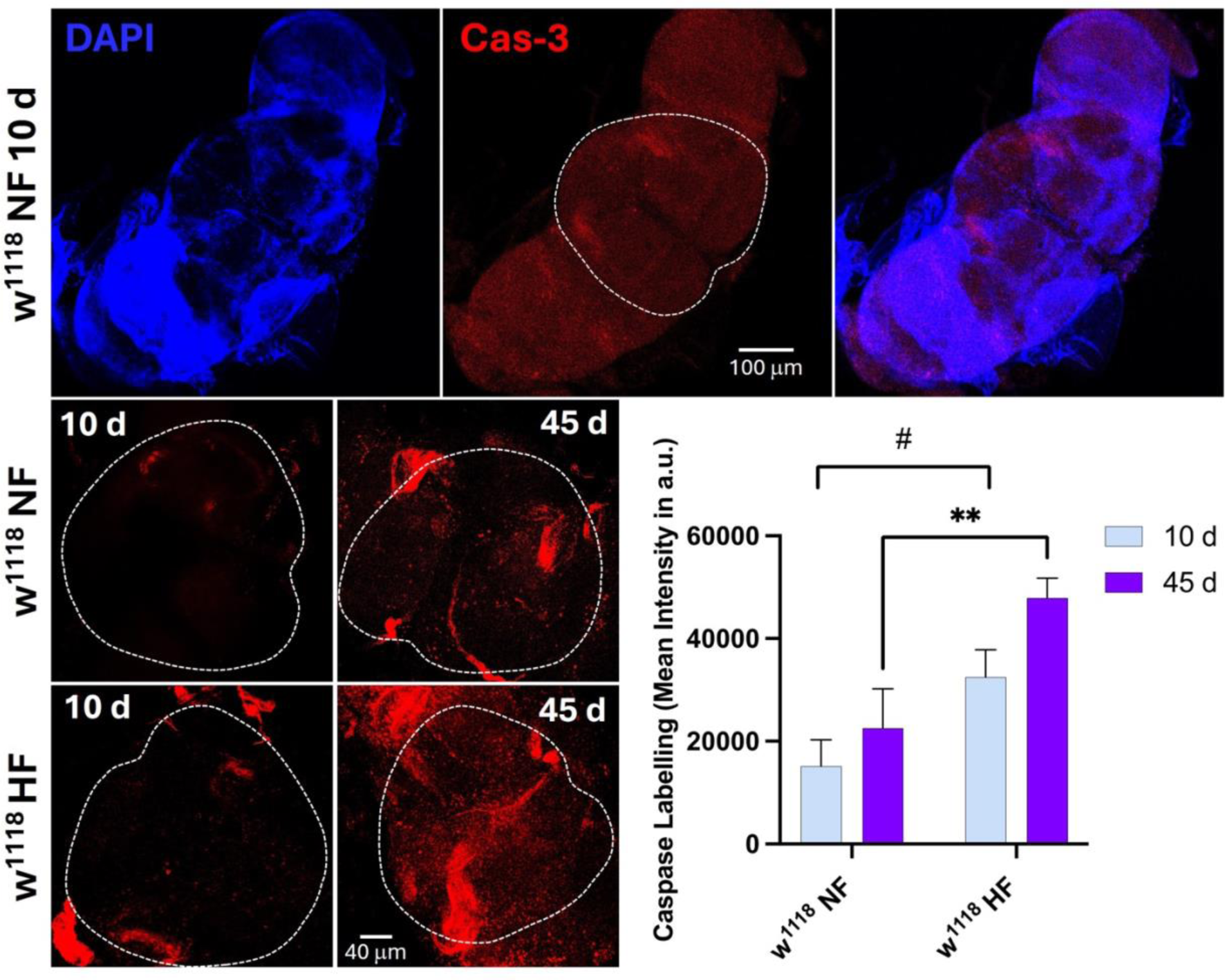
Cleaved caspase-3 levels were enhanced in a diet-dependant manner. A, Representative images of maximal intensity projects at 10 days of NF male (20x); dotted line indicates DAPI-positive inner brain region used for analysis. B, Images of maximal intensity projections of caspase-3 signals were measured in indicated inner brain regions (40x) (diet: F (1, 14) = 12.88, p=0.0030, age: F (1, 14) = 3.688, p=0.0754, NF vs HF at 45 days: **p=0.0093 (two-way ANOVA); NF vs HF at 10 days: ^#^p=0.0271 (unpaired Student’s t-test)) n=6-9 brains per condition.

However, an unpaired Student’s t-test comparison also revealed a caspase-3 signal increase at 10 days following a HF diet (w^1118^ NF: 10 days vs w^1118^ HF: 10 days, p=0.0271, **Figure 6**). Overall, the data suggest an increased apoptotic signalling following a HF diet which would compromise neuronal circuitry function in line with observed behavioural changes reported above.

## Discussion

Our data has revealed new insights into how exposure of *Drosophila* larvae and adult flies to a HF diet exerts differential effects in male and female animals in behaviour and disease marker expression. The main findings illustrate how a HF diet reduces learning and memory function in 3^rd^ instar larvae. In the adult flies, an age-dependent reduction of climbing abilities was detected at younger females at 10 days of age exposed to a HF diet. These behavioral changes are paralleled by gender and age-dependent changes in MDA levels which showed an increase with age (at 45 days of age) in male and female controls but also at 5 days of HF diet (10 days of age). Interestingly at 45 days, MDA levels were lower in both genders, with males exhibiting higher levels at both ages compared to females. An increase in cleaved caspase-3 levels was observed in both sexes exposed to the HF diet although life span was not affected.

A HF diet can exert detrimental effects on brain health and contribute to the pathogenesis of neurodegenerative diseases through neuroinflammation, dysbiosis and dysregulation of the gut-brain axis. By using a *Drosophila* HF diet preparation to explore the mechanism of diet-induced inflammation in remote tissues, a study found that a HF diet (30% coconut oil) activated the NFκB immune pathway in the brain through remodeling of the gut bacteria resulting in increased abundance of *Acetobacter malorum* (Wang, Gu et al. 2024). In addition to dysbiosis, the metabolome of HF diet (20% coconut oil) exposed flies exhibits strong alterations within the first 9 days (Cormier, Doiron et al. 2023). These changes include fundamental metabolic abnormalities such as regulation of the tricarboxylic acid (TCA) cycle and glycolysis, as well as an accumulation of medium/long chain fatty acids. Furthermore, a 2-day HF diet (20% coconut oil) induced mitochondrial fragmentation linked to mTORC2-mediated mitochondrial fission, likely a cytoprotective response to elevated oxidative stress upon HF diet exposure (Liu, Chang et al. 2022). Several studies investigating effects of a HF diet show variable outcomes on different levels of cellular function, however, due to the differences in treatment, coconut oil concentrations *versus* length of exposure, direct comparisons between studies are challenging. The above studies did not separate genders for their analyses which may occlude potential sex-dependent changes.

In order to specifically assess the effects on gender during aging, we analyzed various neuronal phenotypes in our study.

Although there is a general agreement that a HF diet reduces life span using between 20% to 30% coconut oil or 10% lard (Trindade de Paula, Poetini Silva et al. 2016, Wang, Sun et al. 2017, Wen, Zheng et al. 2018, Rivera, McHan et al. 2019, Chattopadhyay and Thirumurugan 2020, Liao, Amcoff et al. 2021, Nayak and Mishra 2021), some studies reported enhanced life spans in fly and rat following a HF diet (7% lard) exposure (Shi, Han et al. 2021). More interestingly, different studies performed longevity experiments under different temperatures: 23°C (Miquel, Lundgren et al. 1976, Wen, Zheng et al. 2018, Rivera, McHan et al. 2019, Molon, Dampc et al. 2020), 25°C (Chattopadhyay and Thirumurugan 2020), 29°C (Zakharenko, Bobrovskikh et al. 2024) which confirms a reduced life span at higher temperatures with greatest effects of a HF diet at higher temperatures. Our data is in agreement with these findings showing largely increased life spans at 18°C although also suggesting a loss of HF diet effects (Figure 4).

In general, in our longevity studies performed at low temperature flies will have a decreased overall metabolic rate, reduced amount of damage of macromolecules (nuclei acids, proteins, lipids) in cells caused by reactive oxygen species (Molon, Dampc et al. 2020) and therefore slowed aging. An additional more general phenomenon of colder acclimatization induces moderate changes in cryoprotective substances and metabolic robustness as shown in *Drosophila suzukii* (Enriquez, Renault et al. 2018) which will slow aging processes and may explain a lack of negative HF diet effects.

Another important finding based on the fact that a HF diet regulates the mRNA or protein expression of genes related to lipid synthesis, including Acetyl-CoA carboxylase and fatty acid synthase (FAS), and genes related to lipid oxidation (Shi, Han et al. 2021) explains life span extensions under HF dietary intake. Interestingly, similar positive effects on life span by downregulating FAS have been shown in *Drosophila* supplemented with a probiotic and synbiotic formulation (Westfall, Lomis et al. 2018). This data further illustrates that some aspects of age and age-related conditions, particularly inflammation and redox stress are preventable with adequate probiotic and prebiotic treatment.

Shi et al (2021) also found that a HF diet (7% lard) significantly reduced palmitic acid levels, which upregulates the expression of Peroxisome Proliferator Activated Receptor Gamma (PPARG), resulting in reduced oxidative stress and inflammation and prolonged lifespan. Activation of this pathway may be able to counteract some of the negative effects of a HF diet seen in our study and may explain the lower MDA levels at 45 days at the HF diet at both genders (Figure 5). Furthermore, our data suggest that apoptotic cell death is enhanced in HF diet exposed flies (Figure 6) most prominently at 45 days but also detectable at 10 days. At this point we have not identified the neurons or circuitries which might be affected by cell death, nor can we suggest that this increase in cleaved caspase-3 signal will have functional consequences affecting climbing activities directly.

The finding that at 10 days of HF diet exposure increased levels of caspase-3 are detectable indicates CNS neuronal dysfunction at this age. Our negative geotaxis data suggest that particular in females, the climbing abilities are compromised at 10 days, suggesting that females may be more susceptible to HF diet insults. Due to the larger n-numbers in this assay, the diet effect was detectable in comparison to the lower n-number and greater variability ICC studies lacking a gender difference.

Our data in larvae suggest a lack of CNS/VNC-motoneuronal effects caused by the HF diet, yet, learning and memory function is impacted. One possible explanation for the differential effects may be caused by differences in redox sensitivity between neuronal populations. Redox stress affects neuronal and synaptic activities and can be mediated via reactive oxygen (ROS) and reactive nitrogen species (RNS) (Wang and Michaelis 2010, Spiers, Breda et al. 2019). A 2% coconut oil HF diet exposure at 24°C (Nayak and Mishra 2021) or 10/20% at 25°C (Trindade de Paula, Poetini Silva et al. 2016) induces ROS which in the presence of neuronally released nitric oxide (NO) as a result of *Drosophila* NO synthase (*d*NOS) activation (Andreakis, D’Aniello et al. 2011) will produce various reactive species involved in pathological regulation of the CNS (Stamler, Simon et al. 1992, Steinert, Chernova et al. 2010, Robinson, Bourgognon et al. 2018, Bourgognon, Spiers et al. 2021). Sensitivity to redox stress has been shown in the case of an accelerated aging of cholinergic neurons within the olfactory circuit in adult flies due to enhanced levels of oxidative stress (Hussain, Pooryasin et al. 2018). Although this was a mixed gender study, overexpression of superoxide dismutase could slow the aging process but no conclusion about sex differences could be drawn. This neuron-specific redox stress sensitivity may explain why *Drosophila* larvae in our study exhibit reduced olfactory learning (Figure 2) but no defects in locomotor activity (Figure 1).

Together, our data show diet-specific changes in behaviour and disease marker expression, with some sex differences in climbing and MDA formation. Future studies will focus on the underlying mechanisms to explain those differences but involving targets such as gut microbiota and gut-derived metabolites represents a potential interventive approach for mitigating the adverse effects of high-fat diets to induce neurodegeneration. Studies should focus on elucidating the neuron-specific redox pathways and microbial and associated molecular mechanisms underlying the gut-brain axis in neurodegenerative disorders.

## Acknowledgements

This work was supported by the University of Nottingham (JRS). We thank Maria Haig for supporting the studies and Ian Ward and the SLIM facility for the continued support of imaging studies.

## Conflict of interest

The authors declare no conflict of interest.

## Author Contributions

ML, VY, KF, CO, RP, IH, MJ, SW performed and analyzed learning and memory, ICC and longevity studies and helped write the manuscript. JRS conceptualized the project, analyzed data and wrote the paper.

## Notes

### Competing Interest Statement

The authors have declared no competing interest.

### Summary of Updates

This version has been formatted with no changes made to the content.

